# A step-by-step sequence-based analysis of virome enrichment protocol for freshwater and sediment samples

**DOI:** 10.1101/2020.09.17.302836

**Authors:** Federica Pinto, Moreno Zolfo, Francesco Beghini, Federica Armanini, Francesco Asnicar, Andrea Silverj, Adriano Boscaini, Nico Salmaso, Nicola Segata

## Abstract

Cultivation-free metagenomic analysis afforded unprecedented details on the diversity, structure and potential functions of microbial communities in different environments. When employed to study the viral fraction of the community that is recalcitrant to cultivation, metagenomics can shed light into the diversity of viruses and their role in natural ecosystems. However, despite the increasing interest in virome metagenomics, methodological issues still hinder the proper interpretation and comparison of results across studies. Virome enrichment experimental protocols are key multi-step processes needed for separating and concentrating the viral fraction from the whole microbial community prior to sequencing. However, there is little information on their efficiency and their potential biases. To fill this gap, we used metagenomic and amplicon sequencing to examine the microbial community composition through the serial filtration and concentration steps commonly used to produce viral-enriched metagenomes. The analyses were performed on water and sediment samples from an Alpine lake. We found that, although the diversity of the retained microbial communities declined progressively during the serial filtration, the final viral fraction contained a large proportion (from 10% to 40%) of non-viral taxa, and that the efficacy of filtration showed biases based on taxonomy. Our results quantified the amount of bacterial genetic material in viromes and highlighted the influence of sample type on the enrichment efficacy. Moreover, since viral-enriched samples contained a significant portion of microbial taxa, computational sequence analysis should account for such biases in the downstream interpretation pipeline.

**Importance:** Filtration is a commonly used method to enrich viral particles in environmental samples. However, there is little information on its efficiency and potential biases on the final result. Using a sequence-based analysis on water and sediment samples, we found that filtration efficacy is dependent on sample type and that the final virome contained a large proportion of non-viral taxa. Our finding stressed the importance of downstream analysis to avoid biased interpretation of data.

## Introduction

Viruses populate all kinds of ecosystems, from natural environments to human-associated ones (e.g. the gut). Their ecological importance derives not only from their astounding abundance - being the most abundant biological entities on Earth (1) - but also from the key role they play within microbial communities. In aquatic systems, viruses can regulate the microbial community influencing biogeochemical cycles and driving the exchange of genes between prokaryotic cells (2, 3). Water in the environment can comprise up to 10^4^-10^8^ viral-like particles (VLP) per millilitre, but such viral diversity is still largely uncharacterised and unexplored (1). Next generation sequencing of environmental genetic material (metagenomics) has allowed the exploration of microbial diversity to an unprecedented detail (4–7). However, some key methodological limitations hinder the quantification of viral diversity and the characterization of their function within the microbial community. The small viral genome sizes that bias in nucleic acid extraction (<1 ng μl^-1^), and the lack of universally conserved genomic regions in viral genomes are common issues faced during virome analysis, particularly for environmental samples of complex matrix such as soil or sediment (8). The separation between viral particles and the solid phase can be difficult because of their strong interactions, which depend on the physico-chemical characteristics of the particulate matter (9, 10). In order to separate viral-like particles (VLP) from particulate matter and microorganisms and to increase virus concentration (and thus viral genetic material), VLP enrichments protocols are employed. Current VLP enrichment protocols use several steps, such as dissolution, centrifugation, filtration and purification/concentration, which can vary from study to study (11–14).

Benchmark investigations have revealed that different viral enrichment protocols could generate different biases on the final virome product, mostly related with microbial contamination and biases against specific viruses (15–19). These findings call for strong caution on profiling and detection of VLPs in viromes, specifically when associations between pathologies and samples/microorganisms are claimed (20).

Filtration is a size-based procedure that is commonly used as a separation step for virome enrichment analysis. Viruses have generally size in diameter between 0.02 μm to 0.4 μm (21, 22). The filter’s pore size of 0.45 μm and/or 0.22 μm are normally adopted, assuming that only particles smaller than their pore size would pass through the filter and that the resulting filtrate would be therefore free of microbial cells, and enriched with viruses. However, several investigations of aquatic ecosystems revealed bacteria able to pass through 0.22 μm filters (23). Presence of microbial genetic material has been broadly confirmed in a recent meta-analysis of viromes studies from human, animal and environmental samples. This highlighted how enrichment protocols can hinder the correct analyses of viral communities because most of the viromes were contaminated by bacterial, archaeal and fungal genetic material (18).

Although studies comparing and optimizing different enrichment protocols have been conducted (15–18, 24), a detailed examination of the efficacy of filtration and the effects at the microbial community level (microbial community composition) is lacking.

The aim of this study was to understand the effect of filtration on virome preparation. In particular, we tested its effectiveness in removing microbial cells from viral-enriched filtrate of particulate-associate and aquatic-based samples. We run a combination of serial filtration and concentration steps commonly used to produce a viral-enriched metagenome. In order to examine the composition of the microbial fraction progressively retained in the filters, amplicon sequencing of DNA recovered at each step was performed paired with shotgun sequencing of viromes. The experiment was performed with sediment and water samples of an Alpine lake, as representative of typical environmental samples.

## Results

To test the effect of multiple filtrations on the composition of the input microbial community and on the induced relative abundance of the viral fraction, we performed multiple consecutive filtration steps on fresh water and sediment samples and sequenced the retained material at each step. Samples of sediment and water were collected at the deepest point (X) and along the coastline (Y) of Lake Caldonazzo, a perialpine lake in Northern Italy. Water was sampled from the epilimnion (WE), thermocline (WT) and hypolimnion (WI). After filtering the input material through three filters of decreasing pores size (10, 5 and 0.22 μm), amplicon 16S rRNA gene sequencing was performed on each filtering to assess the richness and composition of the bacterial component. Genetic material extracted from the raw sediment was also sequenced. However, the DNA retrieved by the extraction of unfiltered lake water was under the detection limit. Therefore, the sequencing was impossible to be applied on such samples. Final enriched samples (viromes) were also analysed by means of shotgun metagenomic sequencing.

The amplicon sequencing data included 33 samples (4 sediment microbiomes, 25 filters and 4 viromes) with a total number of operational taxonomic units (OTUs, clustered at 97% similarity) of 12,104. Shotgun sequencing obtained 322,414,388 quality-filtered paired-end reads, which were on average 92% of the initial reads (**Table 1**). Taxonomic profiling of the shotgun reads with MetaPhlAn and Kraken (25, 26) showed that 99% of the reads in viromes and 60% in sediment metagenomes were not assignable (i.e. “genetic dark matter”). Differences in microbial composition were identified between particulate-associate and aquatic-based samples (Analysis of variance, F=6.24, p<0.01).

**Table 1.**
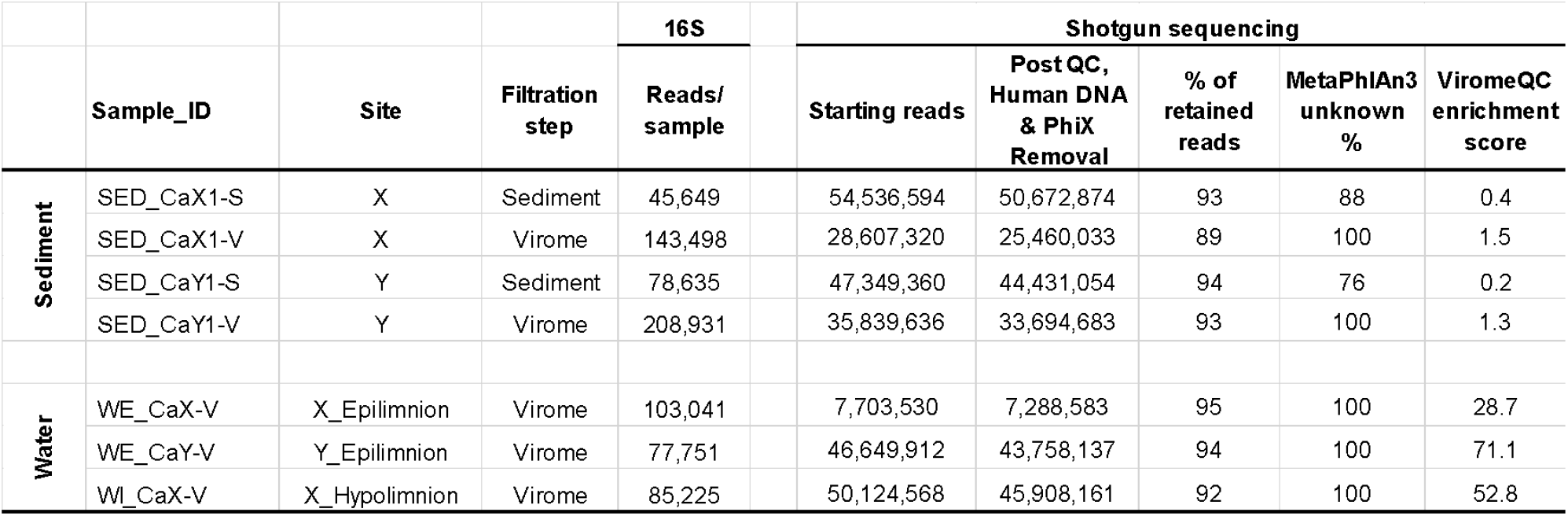
16S and shotgun sequencing reads statistics. SED: sediment, WE: water epilimnion WI: water hypolimnion. X: deepest point of the lake, Y: by the coastline. Extended statistics are reported in Supplementary Table 1.

### Viral enrichment scores are sample-type dependent

We evaluated the amount of bacterial contamination in the enriched samples obtained at the end of the filtration steps. Specifically, we compared the relative abundance of DNA fragments from ribosomal genes (16S/18S rRNA and 23S/28S rRNA genes) and universal bacterial markers (31 markers in total) in the initial unfiltered samples with the final viral enriched filtrates (viromes) using the ViromeQC tool (18). Water and sediment viral enriched filtrates (viromes) contained microbial genetic materials, as shown by the number of OTUs detected by the amplicon sequencing (**Table 1**). We found a modest viral enrichment score for the sediment samples with less than 50% bacterial depletion compared to ViromeQC compendium of water unenriched metagenomes, and an enrichment compared to metagenomes from the sample specimen smaller than one order of magnitude (6.5X for site Y and 3.75X for site X) (**Table 1**).

Filtration performed better in the water samples, with an enrichment score between 28.7X and 71.1X compared to the ViromeQC reference unenriched water samples. Nonetheless, a total of 417 reads (0.001% of the total) still mapped against 16S rRNa or 23S rRNA genes even for the most enriched samples (WE_CaY_V, enrichment score = 71x).

### Effect of filtration on microbial diversity and on the detection of rare taxa

We next sought to examine how the different filtration steps impacted bacterial richness and diversity. While the number of total OTUs at the initial filtration step was higher in sediment compared to water (respectively 3780 ± 1638 and 1058 ± 963, **Fig 1A**), a similar number of OTUs were present in the final enriched samples (1062 ± 243 and 759 ± 79). This implies that the filtration performed differently between sediment and water matrix (72% and 28% decreases in OTUs, **Fig 1A**). In sediment, Shannon diversity decreased along the filtration steps, indicating a progressive elimination of the less abundant bacteria (Kruskal-Wallis, p<0.01). Abundant OTUs were still present at the last step of filtration, hiding the detection of low abundant OTUs (detection level 0.2%) (**Fig. 1D**). Conversely, in water samples, filtration removed bacterial OTUs more homogeneously, with diversity and evenness indices remaining almost stable along the filtration process (Kruskal-Wallis, p>0.05., **Fig 1B-C**).

**Figure 1.**
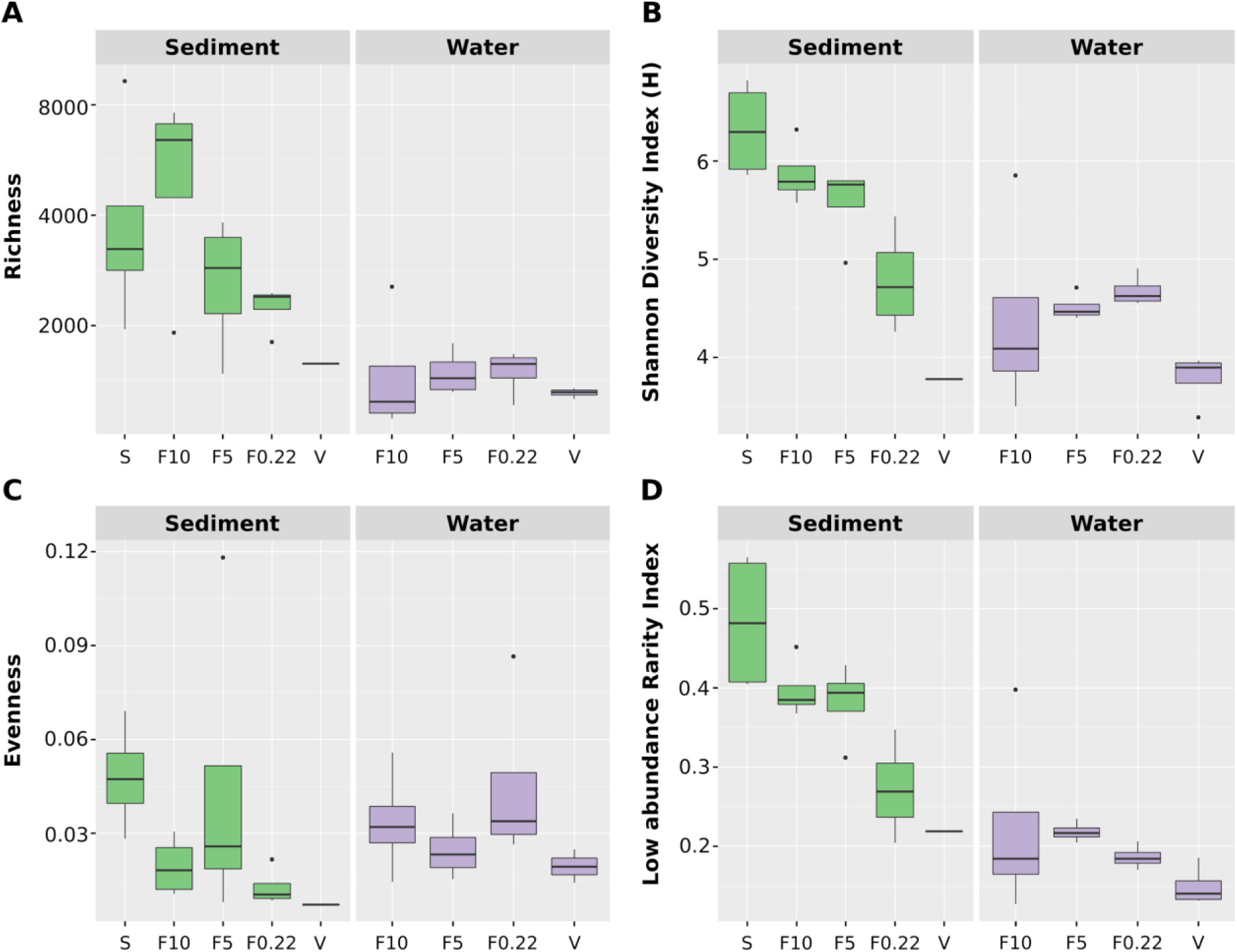
Alpha Diversity indexes of metagenomes, filters and viromes of water and sediments. Boxplots represent the alpha diversity calculated on OTUs from sediments (green) and water (purple). A) Richness, B) Shannon diversity index, C) Evenness and D) Low abundance rarity index. Boxes encompass the quartiles of the distribution, while the median is indicated as a horizontal line in each box. Whiskers extend to show 1.5 Interquartile range. X-axis: filtration categories (S = raw sediment, F10 = filter 10 μm, F5 = filter 5 μm, F022 = filter 0.22 μm, V = virome).

Accordingly, sediment samples displayed a significant decrease of rare species (defined by the rarity low abundance index that measures the relative proportion of species with detection level below 0.2%, regardless of their prevalence) (Kruskal-Wallis, p<0.01. **Fig 2D**), whereas in water samples the relative proportion of rare species remained stable along filtration (**Fig 2D**).

**Figure 2.**
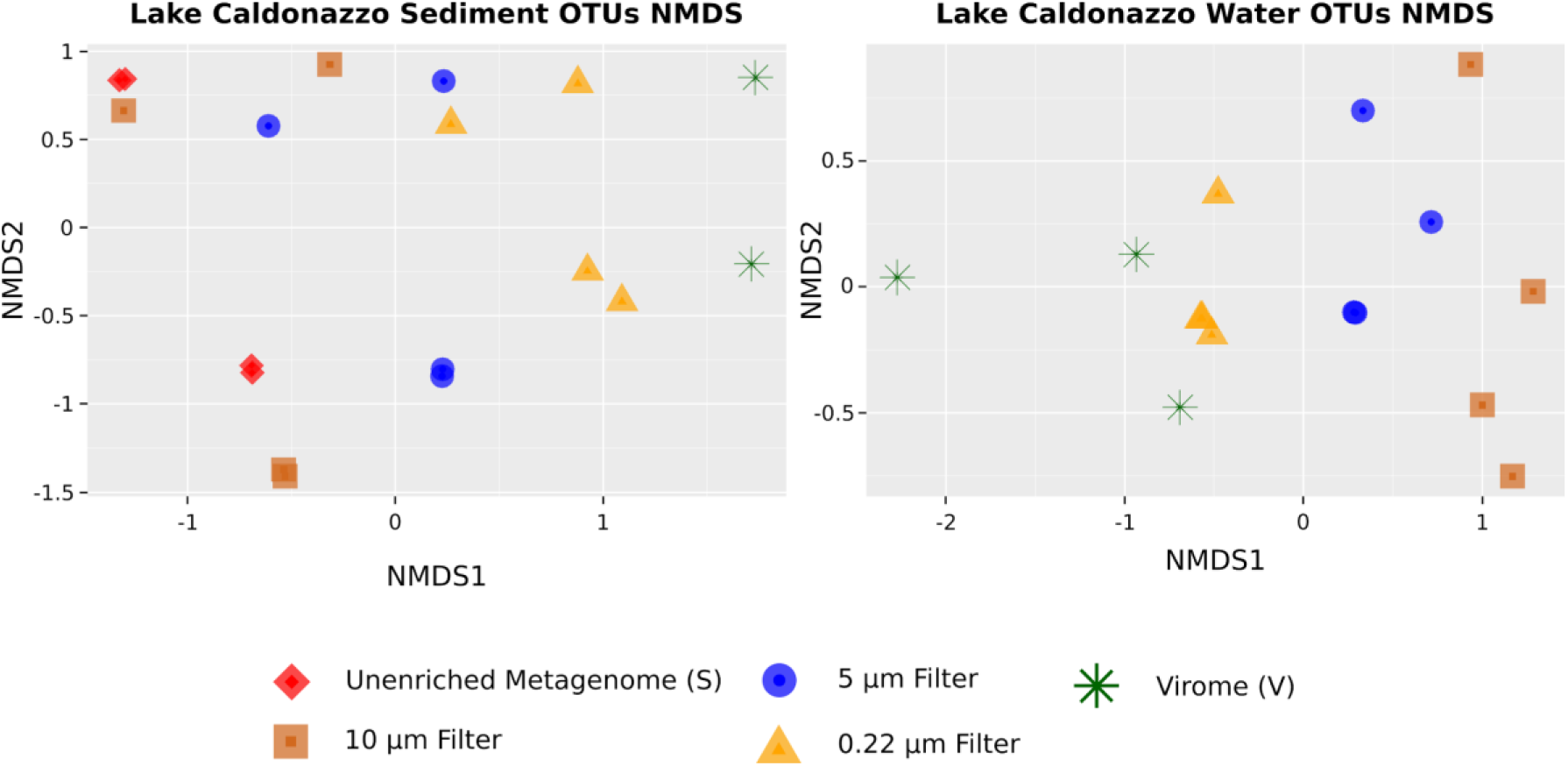
NMDS Analysis on OTUs from metagenomes, filters and viromes of water and sediments. Multidimensional scaling to represent dissimilarity among metagenomes extracted from different pore sizes filters and the viromes of sediment and water samples. OTUs relative abundances were used. Filtration categories: (S = raw sediment, filter 10 μm, filter 5 μm, filter 0.22 μm, V = virome).

### Filtration effects on microbial composition and virome contamination

After assessing bacterial contamination in the enriched samples, we examined bacterial compositional changes induced by filtration steps using 16S rRNA gene amplicon sequencing. As expected, microbial communities differed significantly between water and sediment samples (ADONIS test, p<0.01). Multidimensional scaling (NMDS) of each experimental replicate based on Bray-Curtis dissimilarity showed that samples’ clustering was coherent with filter pore sizes (0.22 μm, 5 μm, 10 μm), which were well sorted along the first NMDS axis both for sediment and water (**Fig. 2**).

We analyzed in greater detail the number of OTUs shared among filters of decreasing pore sizes. Overall, less than one tenth of the OTUs present in the original sediment sample were retrieved after the 0.22 μm filtering step (minimum 2% maximum 9%, **Fig. 3B**) with lower retention rates for the water samples filtration (minimum 11% maximum 28%, **Fig. 3D**). Conversely, the most enriched samples included between 142 (sediment) and 507 (water) unique OTUs that were not detected in the starting (i.e unenriched) samples, and were hence specific of the enriched viromes. While few of these virome-specific OTUs could still be the result of contamination in such low-biomass samples not detected by our computational contamination detection, the majority of these OTUs likely represents taxa that were below the limit of detection in the unenriched samples.

**Figure 3.**
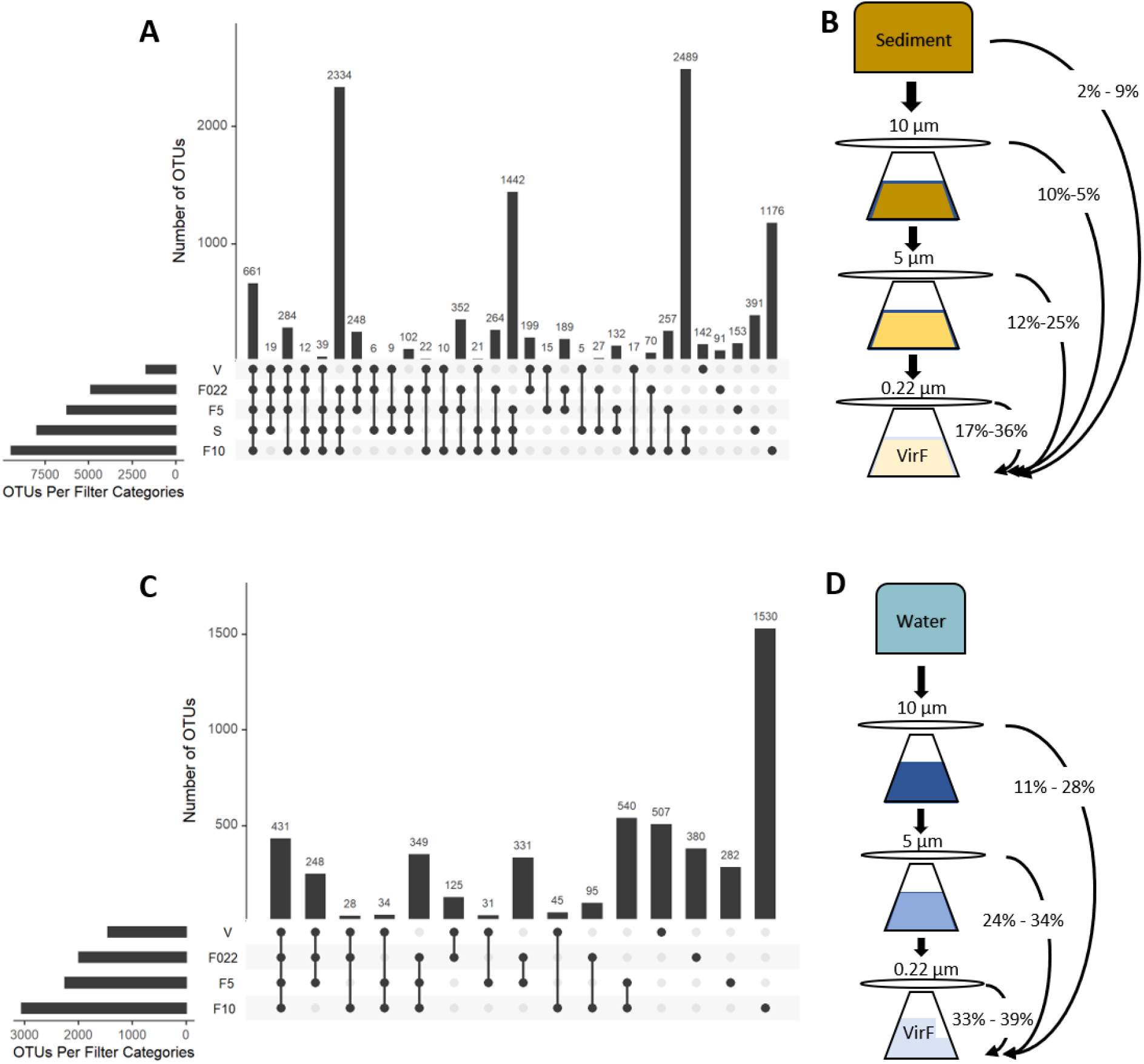
Shared and unique OTUs for the different steps of sequential filtration categories (i.e. the rows). Single dots represent taxa unique to each filtration category; connected dots represent the intersection (shared taxa) among the filtration categories. The y-axis indicates the number of unique or shared OTUs (also reported for each intersection on top of bars) among the categories shown with the dots below. A-B) Sediment samples. A) Shared OTUs among and between filters and viromes. B) Pairwise percentage of shared OTUs, viromes vs filters. C-D) Water samples. C) Shared OTUs among and between filters and viromes. D) Pairwise percentage of shared OTUs, viromes vs filters.

Along the filtration steps, the taxonomic composition of water and sediments differed at phylum (**Fig. 4**) and more evidently at family level (**Supplementary Figure 1**) in the final enriched filtrates. At the initial stage of filtration, Proteobacteria, Cyanobacteria, Bacteroidetes and Planctomycetes were the most abundant phyla. Viromes were still characterised mostly by these phyla, with members of Firmicutes at higher relative abundances (from 5 to 10%). The virome-specific OTUs clearly differed between sediment and water. Sediments were still dominated by Proteobacteria (90%) and Firmicutes (from 5 to 8%), whereas in water a more diverse community was retrieved, including Actinobacteria (from 4 to 8%) and members of the candidate phyla radiation (CPR) (from 0.1 to 3%) such as TM6, Microgenomates, Parcubacteria, Peregrinibacteria, Saccharibacteria and Omnitrophica.

**Figure 4.**
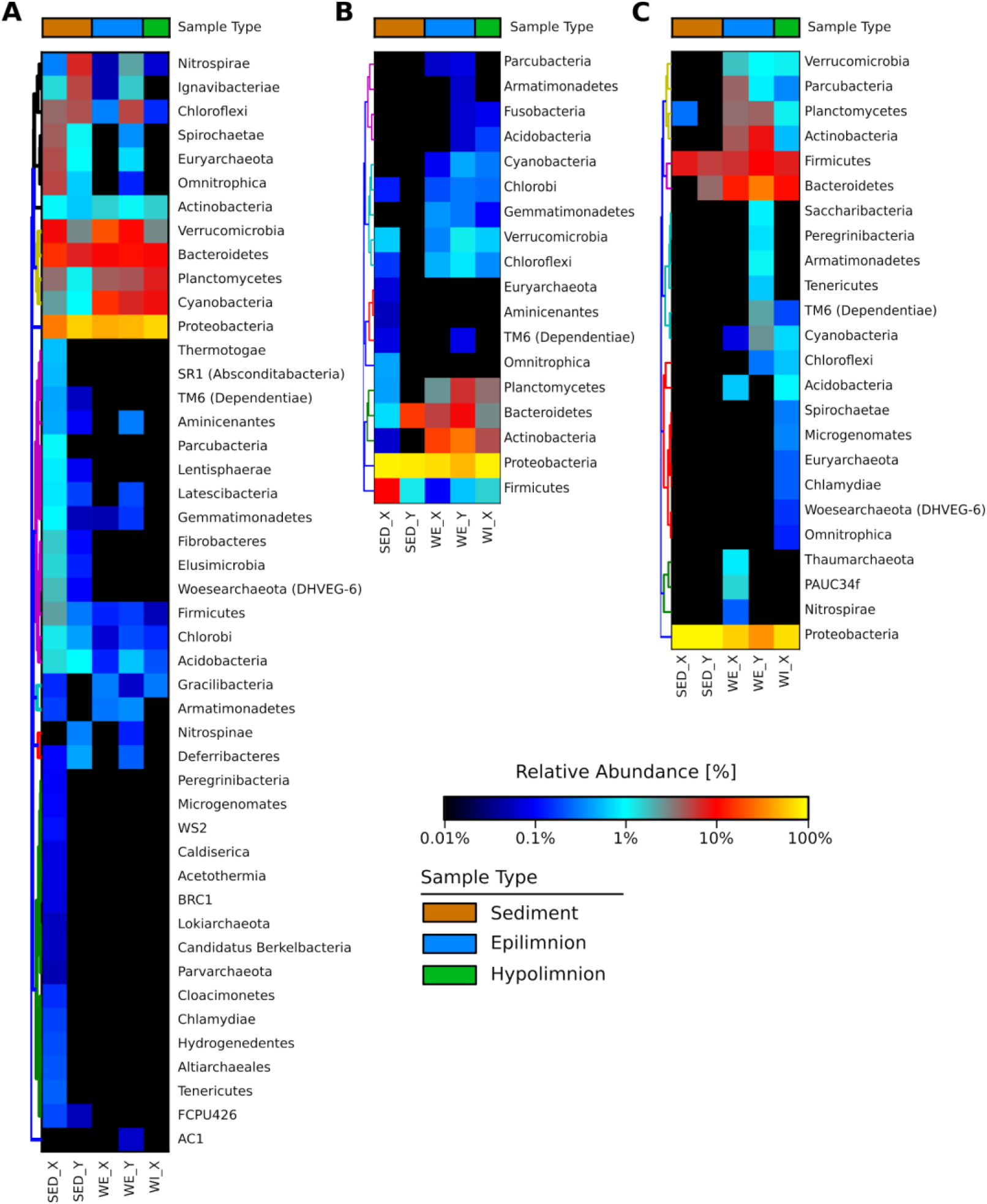
Heatmap of OTUs relative abundance at Phylum level. OTUs relative abundance is shown for A) OTUs at the starting point of filtration. Water samples refer to the OTUs detected in the 10μm filters. B) Enriched viromes total OTUs and C) Enriched viromes unique OTUs. Family level relative abundances are shown in Supplementary Figure 1.

## Discussion

Assigning sequences to viruses poses considerable computational challenges in the study of viromes. The elimination of the genetic material of non-viral origin from the samples is thus fundamental to simplify analyses and avoid biased interpretations. Our study is among the first to specifically test the efficacy of the filtration steps used to eliminate microbial cells from environmental samples during viral enrichment protocols. Particularly, we examined changes in microbial composition occurring throughout the filtration process until the final viral enriched filtrate.

We found that the efficacy of filtration differed between water and sediment samples. The filtration was more effective in water-based samples where it appeared to homogeneously retain microbial species. As such, filtration in water produced viromes that were orders of magnitude more enriched than sediment, as indicated by the ViromeQC enrichment score (18). This result further indicates that the presence of particles and minerals in the sample matrix, such as sediments (27, 28) and, more specifically, sediment characteristics such as porosity and organic matter content, can profoundly influence the efficacy of viral enrichment (12, 29). Consequently, different approaches have been used to account for the retention properties of similar matrices (e.g. soil/sediment, faeces, respiratory samples), such as sample homogenization, as employed here (21, 30, 31). However, our results indicate that additional investigations are needed to develop better laboratory protocols.

Water samples produced much higher enrichment scores, and yet we still detected substantial microbial genetic material in the final enriched samples. This calls for caution when downstream sequence analyses are performed, even after apparent successful enrichment, as it cannot be assumed that the sample contains only viral particles.

Examination of changes in microbial diversity during filtration, provided additional details on how the process differed between sample types. The progressive decline in Shannon diversity and rare species along the filtration steps in sediment samples implies that the efficacy of filtration primarily reflected the relative abundance of taxa, whereby common taxa were more likely to pass through. Conversely, filtration of taxa in water samples appeared to be less dependent on their relative abundance, with both common and rare taxa equally likely to be retained, as indicated by the more stable diversity values. This suggests that filtration in sediments might be relatively more stochastic compared to water samples, where taxa were presumably retained according to their cell size, rather than to their abundance. As previously mentioned, the presence of particle aggregates in the sediment matrix might explain these results, and the lower efficacy of the enrichment.

Although the enrichment differed between sample matrices, filtration steps produced consistent compositional changes across replicate filters in both water and sediment samples, with the first NMDS axis mirroring the distribution of pore sizes (**Fig. 2**). This indicates that, regardless of the overall efficacy, filtration procedures can produce consistent and reproducible outcomes within a given sample matrix.

The key assumption of the enrichment protocols is that only particles smaller than the minimum filter pore size (0.45 μm and/or 0.22 μm) are able to pass through (22). However, in line with other recent studies (17, 18, 32, 33), results from our experiments indicate that viral enriched samples still contained microbial genetic material. This could have practical implications in many research fields. A recent meta-analysis of viromes studies from human, animal and environmental samples, highlighted how commonly used enrichment protocols can hinder the correct analyses of viral communities because of contamination by bacterial, Archaea or fungal genetic material (18). Besides contamination occurring during the experimental procedures, the detection of microbial genetic material in the enriched filtrates could be associated to i) changes in cell size and shape due to external factors; and ii) presence of very small bacteria, such as those belonging to the newly discovered Candidate Phyla Radiation (CPR) (34–36).

Together with the presence of Planococcaceae, Pseudomonadaceae and Sphingobacteriaceae that are commonly found in aquatic and terrestrial habitats (37, 38), viromes also included material from rod-shaped cells such as Oxalobacteraceae, anaerobic purple sulfur Chromatiaceae (39, 40) in sediment and Bryobacter (Acidobacteria) (41) in water. These are small bacteria of 0.3 μm - 0.5 μm cell width, which could pass through the smallest pore size filter (42).

Among the water virome unique OTUs, candidate phyla radiation (CPR) were retrieved. These small bacteria (0.009 ± 0.002 μm) such as TM6, Microgenomates, Parcubacteria, Peregrinibacteria, Saccharibacteria and Omnitrophica, were detected in the enriched final filtrates but were under the limit of detection level at the starting point of filtration. Thus, apparently efficient filtration might enrich not only viral particles but also low abundant microbial species.

Overall, our examination of microbial community diversity and composition associated with the standard virus enrichment protocols, highlight how non-viral particles can be relatively abundant in environmental enriched viromes. We argue that additional effort is needed to further optimise and test viral enrichment approaches, and that researchers analysing and profiling VLPs should be aware of their potential presence.

## Materials and Methods

### Study site and sampling

Caldonazzo Lake is a meso-eutrophic lake located at an elevation of 449 m in Trentino, Italy. Sampling occurred in March 2017 during the lake stratification period in two sites, at the deepest point (X, 49 m depth) and close to the coastline (Y, 7 m depth). Specifically, the first 2 cm of four sediment cores were collected in duplicates and pooled together to collect in total 200 g of sediment. Water samples (2 L) were collected from the two sites (X and Y) at different depths: at the epilimnion (WE. 3 m), thermocline (WT. 10 m) and hypolimnion (WI. 49 m) of the stratified lake. All bottles and devices were acid rinsed and autoclaved before use.

### Microbial and viral DNA extraction

Sediments (100 g) were treated with sodium pyrophosphate (final concentration 5 mM), sonicated and centrifuged in order to separate and collect the sediment pore water (100 mL). Samples (2 L of lake water and 100 mL of sediment pore water) were then serially filtered through 10 μm, 5 μm and 0.22 μm filter pore size (**Fig. 5**) (Whatman filter, Merck KGaA, Darmstadt, Germany) using sterilised filtration units (Nalgene, Thermo Fisher Scientific, USA) mounted on sterile glass bottles. Filters were stored at −20 °C. Virus-like particles (VLPs) in the final filtrate, defined here as the viral fraction, were then concentrated using the iron chloride precipitation protocol (43) and Amicon Ultra filters (100KDa), reaching a final volume of 1-2 mL. Samples were stored at −80 °C.

**Figure 5.**
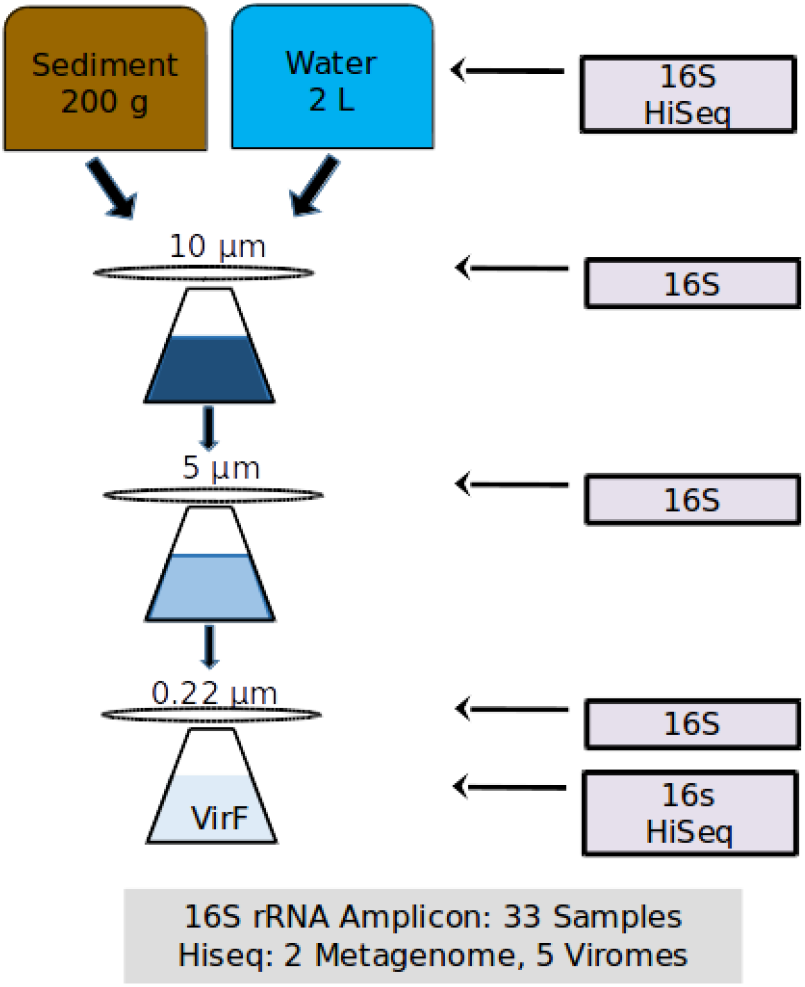
Overview of the extraction procedure. Overall, 33 16S rRNA gene amplicon libraries and 7 shotgun libraries were extracted from freshwater and sediments. The type of library is indicated in the gray boxes on the right. VirF stands for “Viral Fraction”. Pore sizes are indicated above each filter.

DNA was extracted from both filters and viral fractions using different protocols. The 10, 5 and 0.22 μm pore-size filters were processed using the DNeasy PowerWater Kit (QIAGEN, Hilden, Germany) following the manufacture instructions. Viral fractions, instead, were first treated with DNase I (15U mL^-1^) for 1 h at 37C; then DNA was extracted using QIAamp DNA Mini Kit (QIAGEN, Hilden, Germany). Metagenomes DNA were extracted using DNeasy PowerSoil Kit (QIAGEN, Hilden, Germany) directly from sediment (250 mg) following the manufacture instructions. The extraction was also performed with the unfiltered lake water, but the retrieved genetic material was under the detection limit. Therefore, the sequencing was impossible to be applied on such samples. The DNA was quantified using the Qubit™ dsDNA HS Assay Kit (Life Technologies, Carlsbad, CA).

### 16S rRNA Amplicon and shotgun sequencing

To characterise the microbial community along filtration, DNA from filters, from sediments and from viral filtrates were subjected to PCR amplification of the 16S rRNA variable regions V4 (Primer 515f/806r) (44). Amplicons were pooled and sequenced on an Illumina MiSeq platform.

Shotgun sequencing was applied to the DNA extracted directly from sediment (metagenomes) and to viral fractions (viromes). Libraries, prepared using Nextera XT DNA Library Prep Kit (Illumina) according to the manufacturer’s instructions, were quality checked by the Perkin Elmer LabChip GX (Perkin Elmer) and sequenced on a HiSeq 2500 platform (Illumina).

### Bioinformatic and statistical analysis

16S rRNA gene analysis was performed with QIIME with default parameters for demultiplexing, quality filtering, and clustering reads into OTUs (45). Operational taxonomic units (OTUs) were picked with the open-reference approach and the SILVA database release 128 at 97% clustering (46). In R, data were processed using phyloseq (47), vegan (48) and UpsetR (49) packages. Archaea, Chloroplast and Mitochondria were removed from the dataset. For the non-metric multidimensional scaling (NMDS, default square-root and Wisconsin double standardisation of values) community analysis, Bray-Curtis dissimilarity was used after removing rare OTUs (<5 occurrences). From vegan package, adonis analysis was performed to determine the differences between habitats, filters and sampling location. Differences in bacterial diversity indexes over the filtration process were tested using a linear regression model, setting as base level the first step of filtration (raw sediment and filter 10 μm for water). Bacterial richness was log transformed. To determine and represent shared OTUs among categories (filters and viromes) and unique OTUs, upsetR was applied.

Raw metagenomic reads were preprocessed with Trim Galore (50) to remove low quality (i.e. Phred score < 20) and short (i.e. length < 75 bp) reads (parameters: --stringency 5 --length 75 --quality 20 --max_n 2 --trim-n). Metagenomes were analyzed with MetaPhlAn (25) v. 3.0 with the --unknown_estimation option and Kraken2 (26), version version 2.0.8 and Braken (51) To quantify also the percentage of reads that could not be assigned to any taxa, the percentage of “unknown reads” was taken from the output of the two tools (i.e. --unkown_estimation in MetaPhlAn). These percentages are reported in **Supplementary Table 1**.

Viral enrichment was calculated with ViromeQC, a computational tool that estimates the efficacy of VLP enrichment by quantifying the abundance of unwanted microbial contaminants. ViromeQC estimates the abundance of contaminants from the raw metagenomic reads via the 16S/18S and 23S/28S rRNA gene abundances, and from 31 single-copy bacterial markers. ViromeQC version 1.0 (18) was run on the metagenomic reads with the --environmental option.

## Data Availability

The raw sequencing reads of the 16S rRNA amplicon sequencing and shotgun metagenomics were submitted to the NCBI-SRA archive and are available under the BioProject PRJNA658338.

## Acknowledgements

This project received funding from the European Union’s Horizon 2020 research and innovation programme under the Marie Skłodowska-Curie grant agreement No. 704603 to FP and the European Research Council (ERC-STG project MetaPG No. 716575) to NS. We would like to thank Stefano Larsen and Nicolai Karcher for their feedback and technical support.

